# Bumetanide prevents brain trauma-induced depressive-like behavior

**DOI:** 10.1101/443739

**Authors:** Goubert Emmanuelle, Altvater Marc, Rovira Marie-Noelle, Khalilov Ilgam, Mazzarino Morgane, Sebastiani Anne, Schaefer Michael, Rivera Claudio, Pellegrino Christophe

## Abstract

Brain trauma triggers a cascade of deleterious events leading to enhanced incidence of drug resistant epilepsies, depression and cognitive dysfunctions. The underlying mechanisms leading to these alterations are poorly understood and treatment that attenuates those sequels not available. Using controlled-cortical impact (CCI) as experimental model of brain trauma in adult mouse we found a strong suppressive effect of the sodium-potassium-chloride importer (NKCC1) specific antagonist bumetanide on appearance of depression-like behavior. We demonstrate that this alteration in behavior is associated with a block of CCI-induced decrease in parvalbumin-positive interneurons and impairment of post-traumatic secondary neurogenesis within the dentate gyrus of the hippocampus. The mechanism mediating the effect of bumetanide involves early transient changes in expression of chloride regulatory proteins and qualitative changes in GABA(A) mediated transmission after brain trauma. This work opens new perspectives in the early treatment of human post-traumatic induced depression. Our results strongly suggest that bumetanide might constitute an efficient prophylactic treatment to reduce neurological and psychiatric consequences of brain trauma.

## Introduction

Brain trauma is the main cause of disability over the world with a very high prevalence in developed countries (Bondi et al., 2015; Meyer et al., 2008). According to the World Health Organization and the Centers for Disease Control and Prevention (Meyer et al., 2008), brain trauma classification is based on multiple factors such as altered neurological functions, brain area of interest and genetic variations. Altogether these factors lead to highly individualized injuries. Sequels of trauma include low prevalence post-traumatic epilepsies (PTE) with a severity and occurrence depending on trauma severity (Bragin et al., 2016; Kelly et al., 2015), cognitive dysfunctions and depression-like phenotypes are as well often associated (Peeters et al., 2015; Perry et al., 2015; Stein et al., 2015). Following brain trauma, neuronal cell death occurs and more particularly within the neurons of the dentate gyrus of the hippocampus (Ren et al., 2015; Samuels et al., 2015) leading to hippocampal volume reduction (Anacker and Hen, 2017; Samuels et al., 2015). These observations could be related to changes in post-traumatic neurogenesis in the hippocampus. This has been proposed to be a useful marker as an indicator of therapeutic treatment efficacy (Alvarez et al., 2016; Brandon and McKay, 2015).

In a wide range of neurological and psychiatric disorders, GABAergic signaling is affected. Particularly mediated by chloride homeostasis impairment triggered by a down regulation of the main neuronal-specific chloride and potassium extruder, KCC2 and up regulation of the chloride importer NKCC1, respectively (Medina et al., 2014). Similar changes in GABAergic transmission have been reported in different model of brain trauma (Ben-Ari, 2017). This leads to depolarizing and also excitatory action of GABA that could perturb the generation of behaviorally relevant oscillations and integrative properties of brain networks (Ben-Ari, 2017; Kahle et al., 2013; Luscher et al., 2011; Medina et al., 2014; Rivera et al., 1999). These shifts have been observed notably in developmental disorders including Autism Spectrum Disorders (ASD) (Tyzio et al., 2014), stroke (Jaenisch et al., 2010; Xu et al., 2016) and epilepsy (Kelley et al., 2016; Pallud et al., 2014; Tyzio et al., 2014).

Interestingly, the NKCC1 chloride importer antagonist bumetanide has been shown to attenuate many of these disorders like ASD, Parkinson, schizophrenia as well as some CCI-induced squeals. This stresses the therapeutic potential of restoring low (Cl^-^)_i_ levels and an efficient GABAergic inhibition (Ben-Ari, 2017; Damier et al., 2016; Lemonnier et al., 2016; 2013; Xu et al., 2016).

The connection between Major Depressive Disorders (MDD) and GABAergic neurotransmission has been suggested in genetic models of GABA(B)-R knock out (Mombereau et al., 2005) and in studies showing antidepressant effect of potent and selective blockage of GABA(A) transmission (Rudolph and Knoflach, 2011) at both the hippocampus (Boldrini et al., 2013) and mesolimbic system (Kandratavicius et al., 2014). In addition, several observations link chloride homeostasis to secondary neurogenesis through GABA(A) neurotransmission (Luscher et al., 2011; Ostroumov et al., 2016). The generation of new neurons within the DG requires different steps, first the transition from quiescent to proliferative progenitors then their differentiation to immature neurons in a GABAergic-dependent manner (Chell and Frisén, 2012; Moss and Toni, 2013). It’s well accepted that brain trauma alters neurogenesis (Perry et al., 2015; Stein et al., 2015). Within this context, parvalbumin-containing interneurons are potential candidates to explain CCI induced dysregulations as they are both susceptible to death and have a strong role in hypersynchronicity of hippocampal networks (Curia et al., 2008; Drexel et al., 2011; Shiri et al., 2014). In addition, it is accepted that the activity of this class of interneurons could act on secondary neurogenesis by providing a source of ambient GABA (Butler et al., 2016; Hu et al., 2017; Pérez-Domínguez et al., 2017; Song et al., 2012) but little is known about the connection between parvalbumin-containing interneurons and the establishment of post-traumatic depression (Earnheart et al., 2007; Fenton, 2015; Luscher et al., 2011). Moreover, in human depression, their action is far from established (Khundakar et al., 2011; Pehrson and Sanchez, 2015; Smiley et al., 2016).

Although it has been previously shown that bumetanide could have various positive effects in TBI models (Hui et al., 2016; Zhang et al., 2017) and could as well act on secondary neurogenesis in stroke condition (Xu et al., 2016), nothing is known about the early action of this compound prior the establishment of depressive-like behaviors (DLB). Our results showed that brain trauma disrupts chloride homeostasis leading to hippocampal network disturbance and impaired neurogenesis associated with DLB. Early restoration of chloride homeostasis using the NKCC1 inhibitor bumetanide rapidly after trauma attenuates the severity of post-traumatic alterations notably by reducing interneuron apoptosis. This taken together suggests a therapeutic potential of this FDA-approved compound after trauma.

## Material and methods

The French ethical approved all experimental procedures (N°: APAFIS#2797- 2015112016427629v8). All experiment were performed in blind.

### Controlled-cortical impact model (CCI)

10-weeks old C57bl6-J males were housed individually in an enriched environment, maintained in a 12 h light/12 h dark cycle environment with controlled temperature (23 ± 2°C), food and water were given ad libitum. The controlled cortical impact (CCI) procedure was performed using aseptic technique. 30 min before surgery, buprenorphine (0.03 mg/kg) was given intra-peritoneally (i.p). Anesthesia induction is done using 4% isoflurane mixed with air and enriched with oxygen (0.3 %), for the procedure isoflurane is set to 2-2.5% before animals are positioned in a stereotaxic frame (David Kopf Instruments). Body temperature is maintained at 37 ± 2°C with a heating pad (Harvard Apparatus). The impact is done on the right cortex within the boundaries of the bregma and lambda after a craniotomy is done, using a leica impactor (tip diameter 3mm, 6 m/sec speed, 1.5 mm depth and 200 msec duration). Sham animals receive the complete surgery without the impact. Before the experiment, animals were randomly assigned to each group e.g sham-vehicle, sham-bumetanide, CCI-vehicle and CCI-bumetanide. i.p Bumetanide injections were performed twice daily during one week at 2 mg/kg.

### Western blot analysis

Tissues were proceeded as previously described (Kourdougli et al., 2017) and membranes stained using NKCC1 (DHSB, 1:2000) and KCC2 (non-commercial, (Ludwig et al., 2003); 1:5000) then normalized using the corresponding tubulin antibody (Alpha, Covance, 1:10000 or beta, Thermofisher, 1:3000). The relative expression levels are normalized to sham condition.

### Immunohistochemistry

Mice were transcardially perfused with 4% paraformaldehyde then 60μm coronal sections were made and stained overnight at 4°C using KCC2 (1:3000; (Ludwig et al., 2003)) BrdU (Dako M0744, 1:100), DCX (Abcam, 1:1000) and parvalbumin (Sigma P3088, 1:500), The Alexa Fluor-conjugated secondary antibodies (1/500, Invitrogen) used 2 h at room temperature and slices were finally counterstained with Hoechst 33258 (10 μg/ml, Sigma-Aldrich 861405). Images were taken using a confocal microscope with 10x, 20x, 40x or 63x objectives.

### KCC2 subcellular localization analysis

The measure of the distribution of KCC2 fluorescence associated with cytosolic regions, in sham and cci condition, was performed at high magnification (x63 objective) using the Image J software. Plot Profile values were done using the same straight line length applied from the external cell compartment to the nucleus.

### BrdU injections and staining

I.p injection of a 1 mg BrdU solution (Sigma, 10 mg/ml) were performed at 6 days and 1 month post brain trauma to label dividing cells in the S-phase. Mice received two BrdU injections (morning and evening) the day before brain collection. Immunohistochemistry is done using a mouse-BrdU antibody (1:100, Dako). The total number of BrdU-immunopositive cells per section were counted within the granular layer of the dentate gyrus (DG) using an apotome module (BX 40 Olympus).

### Behavioral studies

Animals were habituated to the testing room 24 h before testing. For the open field test, mice were allowed to freely explore the space for 10 min (Noldus apparatus, 38.5 cm x 38.5 cm). Parameters were detected using the Ethovision software (Noldus). The forced swim test paradigm was performed in a 25°C water; the procedure made of 2 min habituation period followed by 4 min recording. Immobility time was manually assessed. Stabilization movements were not counted as swimming movements. The tail suspension test was performed on a 6 min trial and the immobility time was manually counted. The splash test consists in spraying a 10%-sucrose solution to the fur of the animal then animals are video monitored for a 5 minutes period during which latency to first complete sequence of grooming and total grooming time is measured. For the novel object recognition test animals are exposed to an open field arena (38,5cm x 38,5cm x 38,5cm) for a 3-min habituation time, then exposed 3 min with two new objects, finally the last “-min session is done 1hour after with one new object. Time spent close to the new versus familiar object is plotted as a ratio.

### Acute slices preparation

Animals were collected on the first post-traumatic week. After cervical dislocation, the brain was rapidly removed, the hippocampi dissected, and transverse 350 to 450μm thickness slices done using a Leica VT1000S tissue slicer as previously described (Khalilov et al., 2005). Slices were then transferred at room temperature for 1-2 h before chloride and electrophysiological recordings in oxygenated (95 % O2 and 5 % CO2) normal artificial CSF (ACSF)(Khalilov et al., 2005).

### In vitro electrophysiological recordings

Hippocampal slices were individually transferred to a recording chamber maintained at 30–32°C and continuously perfused (2 mL/min) with oxygenated normal or adapted ACSF. Extracellular field recordings were made in using tungsten wire electrodes as previously described (Khalilov et al., 2005)

### Volumetry Analysis

Coronal 10μm thick cryostat sections were stained by cresyl violet, digitized and analyzed using 1.25x objective and computer image analysis system (Optimas 6.51, Optimas Corporation, Bothell, WA). Lesion volume measurement was performed essentially as described (Schaible and Straub, 2014).

### Parvalbumin-containing interneurons quantification

40μm sections were stained using a mouse-Parvalbumin antibody (Sigma-Aldrich, 1:500) and counterstained with Hoechst 33258 (Sigma-Aldrich, 10 μg/ml in PBS). The quantification was done in granular layer of the DG at 40X objective. All experiments were manually performed in blind.

### Statistical analysis

All mean values are given with the standard error mean (sem). Normality was tested for each distribution and was set to 5 %. Two-tailed Student’s, Mann-whitney test or One-Way Anova were used accordingly using Prism software (GraphPad software, Inc., La Jolla, CA, USA). Box plot report the median, the interquartile range and the total range data and represent as following: * p <0.05; ** p <0.01 and *** p <0.001.

## Results

### Behavioral analysis of depressive-like behavior

CCI protocols trigger the appearance of comorbidity factors at later stages e.g. depression-like behavior. We first performed behavioral tests to ensure that the mice model of brain trauma in this study exhibited DLB. We performed forced swim test (FST) for 4 minutes after 2 minutes habituation (Poleszak et al., 2006; Tao et al., 2016), tail suspension test (TST) for 6 minutes (Castagné et al., 2011; Fan et al., 2016), open field test (OFT) for 10 minutes (Tao et al., 2016), splash test for 5 minutes (Marrocco et al., 2014; Petit et al., 2014) and finally novel object recognition (Egeland et al., 2017) in a sequence of 3 min habituation, 3 min exposure to objects and then 1 hour after, 3 min exposure to familiar and novel object (García-Pardo et al., 2016). All those experiments were carried out one month after the CCI. In the OFT we observed significant changes in the time spent by the animal in the center of the arena (sham 50.83 +/- 13 sec vs CCI 60.86 +/- 22 sec, sham n=30, CCI n=20, p=0.04, Fig. 1A) whereas there was no significant difference neither in the total distance (sham 3308 +/- 160 cm versus CCI 3571 +/- 155 cm, p=0,2, Fig. 1A) nor in the average speed of the animals (sham 5.8 +/- 0.2 m/sec versus CCI 6.2 +/- 0.3 m/sec, p=0.3, Fig. 1A). Then we moved to more specific test for depression using the FST and TST paradigm and found a significant increase in the immobility time of CCI animals versus sham (152.8 +/- 12 sec versus 88.8 +/- 6 sec, p<0.0001, n= 12 and 15 animals respectively; Fig. 1B) (165+/- 54 sec versus 246+/- 36 sec, p=0.007; Fig. 1C). To confirm that this was indeed a DLB, we injected imipramine (30 mg/kg, n=20 animals) a classical anti-depressant compound (Castagné et al., 2011; Cryan et al., 2005; Zhao et al., 2015) given 30 min before the test. In agreement with the literature we observed a strong effect on the phenotype (88.8 +/- 6 sec versus 89.6 +/- 30 sec for imipramine treated animals, 6 animals). The same effect on immobility time was observed on the TST (168,6 +/-31 sec versus 246,7 +/- 18 sec, n=10 per condition)(Fig. 1C). Imipramine had a similar effect that the one observed on the TST. The DLB was completed by performing novel object recognition test, a well-known test for MDD (Egeland et al., 2017), here we observed a significant change in the time spent by the animal around the new object after CCI, again this phenotype was rescued by bumetanide treatment (1 +/- 0,13 versus 0,5 +/- 0,07; n=17 per condition) (Fig. 1D). Finally, the grooming of the animals was assessed after a sucrose solution was sprayed to the fur. We observed an increase in the time of the first complete grooming sequence (93,5 +/- 5 sec versus 129,9 +/- 12, n=26 per condition) without any significant change in the total grooming time (168 +/- 7 versus 142 +/- 9, n= 26 per condition) (Fig. 1E).

**Figure 1:**
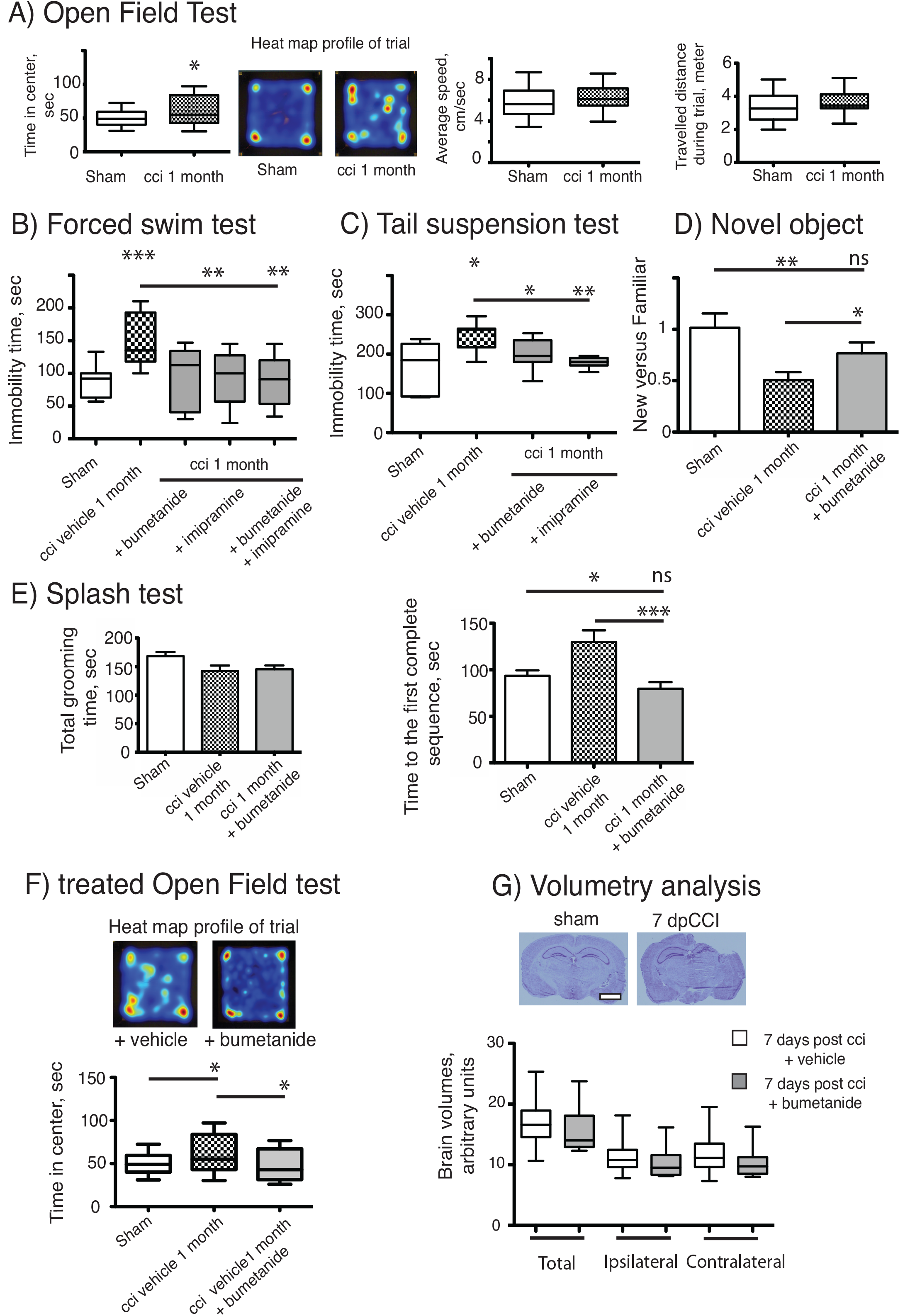
Bumetanide ameliorates CCI induced behavioral changes. A) Open field test: Plots represent both the time spent by the animal in the arena center during the 10minutes test, the total distance travelled and the average speed of the animal during the trial, sham n=30 and CCI n=20. B) Forced swim test: Immobility time in a 25°C water for 4 min, sham n=15, CCI n=12, CCI + bum=6, CCI + imipramine=6 and CCI + bum + imipramine=6. C) Tail suspension test: Immobility time for 6 min (sham n=10 and CCI n=10. D). The Novel Objet Recognition shows changes in the exploration time, the results are presented on a ratio of time of new versus familiar, n=16, 15 and 17 respectively. E). Splash test analyzes the total grooming time and the latency to the first complete sequence. n= 26, 27, 28 respectively. F) After one week of i.p bumetanide injection (20 mM) injection. Treated. Animals showed improvement in the open field test compared to non-treated animals, n=20. G). Volumetry analysis: Volume was calculated by summation of areas multiplied by distance between sections (500 μm) n=10 per condition. The graph shows the mean volume of brain hemispheres, ipsi- and contralateral, 7 days post CCI from bumetanide- and vehicle-treated mice.

### Early application of bumetanide rescues CCI induced depressive-like behavior

We then subjected CCI mice to i.p injections of bumetanide (2 mg/kg) twice daily during the first week after CCI. This time window was chosen as the blood brain barrier is considered to remain open (Dachir et al., 2014). Behavioral analyses of the cohorts were performed again after 1 month. Analysis of these results revealed a potent action of bumetanide on all the behavioral tests. This indicates a major role of CCI induced changes in chloride homeostasis in the induction of DLB. The effect of bumetanide was significant in both on the immobility time using FST (p=0.0002, Fig. 1B) and the TST (p=0,03, Fig. 1C) but also on the exploratory paradigm of the OFT (p=0,04, Fig. 1F), as well as on the splash test (p=0,0005, Fig. 1E) and the novel object recognition test (p=0,0005, Fig. 1D). Interestingly, we did not observe any significant difference in the FST paradigm between imipramine treated animal and bumetanide-imipramine double treated animals (Fig. 1B), indicating no additional effect of bumetanide over imipramine.

The accumulation of intracranial pressure is a comorbidity of close TBI. This is primarily produced by the formation of edema. Previous work suggest that changes in chloride homeostasis can have an ameliorating affect on trauma induced edema (Lu et al., 2007) that could mediate the positive effects of inhibiting changes in chloride transport. Significant formation of intracranial pressure is not expected in this study, as it is an open CCI model.

To investigate the effect of bumetanide on edema, we performed volumetry analysis of the lesion size on both bumetanide-treated and vehicle-treated brains. By cresyl-violet staining, we defined the lesion size in both treated- and non-treated conditions (Fig. 1G). Our results show that at 7dpCCI, there was no significant change in the lesion size neither of the global brain volume (n=10, Fig. 1G) nor on hemispheres (n=10, Fig. 1G). This is in agreement with a previous publication (Lu et al., 2007).

### CCI induced changes in hippocampal network activity and inhibitory strength of GABAergic signaling

Hippocampus is one of the main regions known to be involved in the occurrence of DLB. Studying changes in neuronal activity is important to understand the appearance of MDD. The action of bumetanide on the prevention of post-traumatic DLB suggests that changes in chloride homeostasis and in GABAergic neurotransmission in the hippocampus may be involved in the process. Indeed, impairment in chloride homeostasis after TBI has already been shown in the hippocampus (Rivera et al., 2004; Shulga et al., 2012). In order to asses whether GABAA transmission is affected in the model used in this study we monitored the effect of GABA(A) receptor activation on extracellular field potentials in the hippocampus. As spontaneous activity in DG is known to be quite low (Kvajo et al., 2011; Spruston and Johnston, 1992) (Supplemental figure 1) we decided to record multi-unit activity (MUA) from the CA3 hippocampal region. Acute hippocampal slices both from ipsi- and contralesional hemispheres at 3 days after CCI were recorded and 10 μM isoguvacine, a potent and selective GABA(A) agonist, applied. Such treatment exert an excitatory action on the action potential spiking frequency on ipsi- but not on contralateral hippocampus at 3 dpCCI (Fig. 2B) as compared to sham condition (Fig 2A). This set of results suggests that GABAergic transmission is largely modified making the network more excitable. The strong block of the depolarizing effect of isoguvacine by 10μM bumetanide (n=2 animals, 4 to 5 slices per animal) (Fig. 2C) indicates the involvement of chloride imbalance in the CCI induced changes in GABA(A) responses.

**Figure 2:**
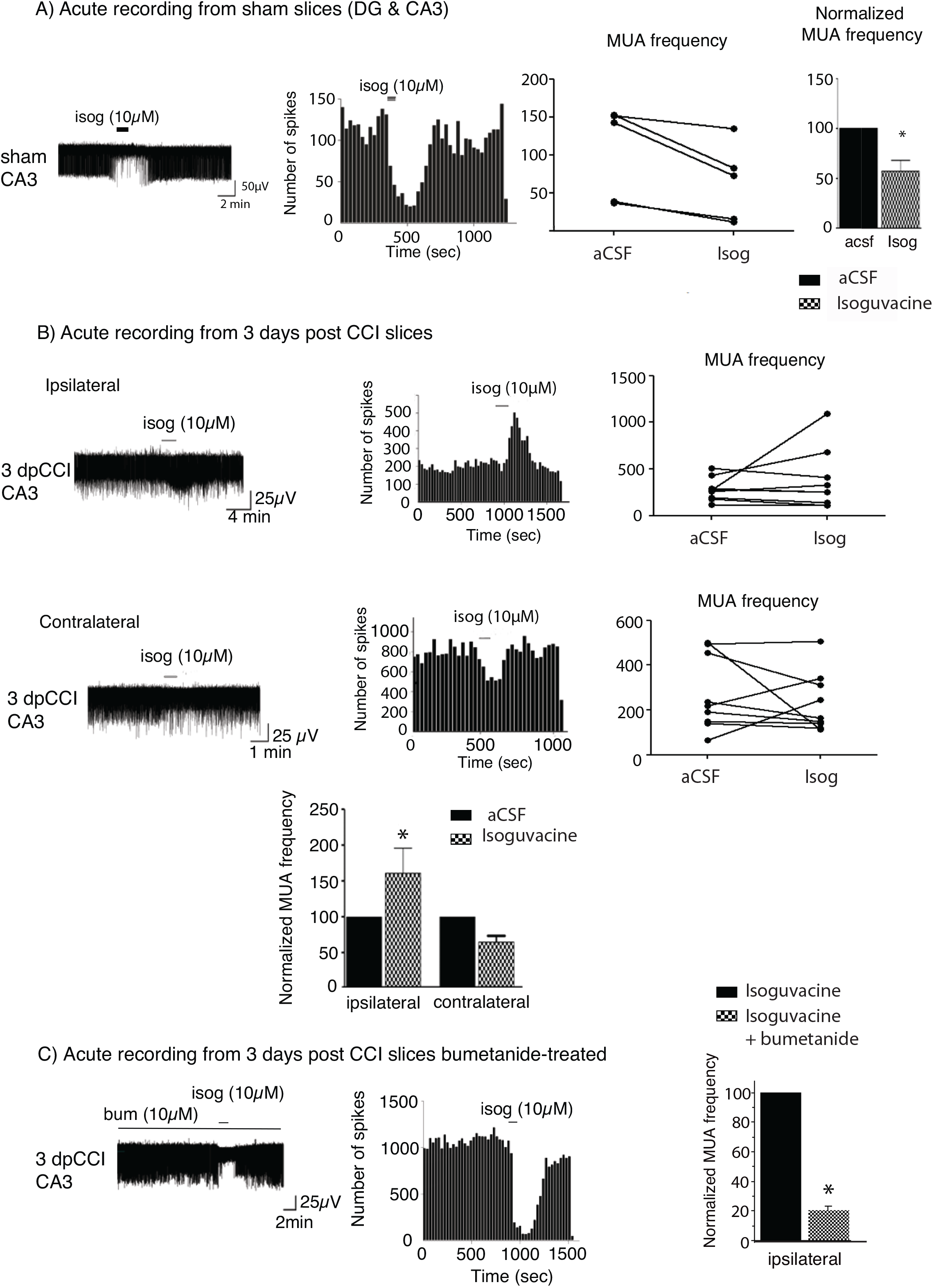
Network activity recording and chloride extrusion efficacy. A) Effect of isoguvacine (10 μM) on hippocampal networks from ipsi and contralateral hippocampus from sham animals. B) Effect of isoguvacine (10 μM) on hippocampal networks at 3 days post CCI, Top left: example trace of spontaneous extracellular field potentials recorded in ipsilateral hippocampus. Middle: corresponding time course of spike frequency changes. Right: graph of non-normalized spike frequencies. Middle left: example trace of spontaneous extracellular field potentials recorded in contralateral hippocampus. Middle: corresponding time course of spike frequency changes. Right: graph of non-normalized spike frequencies. Bottom: average histograms of normalized spike frequencies. C) The same as in (B) with acute pre-treatment of bumetanide (10 μM). 3 days post CCI (n=2 animals, 4-5 slices per animal).

In order to asses the impact of CCI in other regions we decided to measure the functionality of chloride transport using Clomeleon mice (Berglund et al., 2006) (supplemental Figure 2). We found that chloride extrusion was significantly reduced at 3 and 5 but not at 7 dpCCI compared to sham condition. The chloride extrusion capacity of KCC2 is modified and tightly linked to changes in network excitability as observed using MUA recordings. This also suggests a general effect of CCI on chloride transport over the hippocampus.

### Changes in chloride regulatory proteins after CCI

We investigated whether changes in network excitability could be explained by changes in dynamics of chloride extrusion efficacy. To estimate to what extend chloride regulatory proteins are affected during the early post-traumatic time window following CCI we follow expression of NKCC1 and KCC2 during the first post-traumatic week in the hippocampus. We observed a significant decrease in KCC2 protein expression rapidly after the trauma with a recovery at the 7^th^ day at the ipsi and contralateral hippocampus (Fig. 3A&B). For NKCC1 analysis, we did not observe any significant changes in hippocampi from ipsi- and contralateral side (Fig. 3A&B). Similar results are obtained for KCC2 and conversely for NKCC1 at mRNA levels in the injured hemisphere (Supplemental Fig 3). These results show an imbalance in NKCC1/KCC2 ratio in favor of NKCC1 expression.

**Figure 3:**
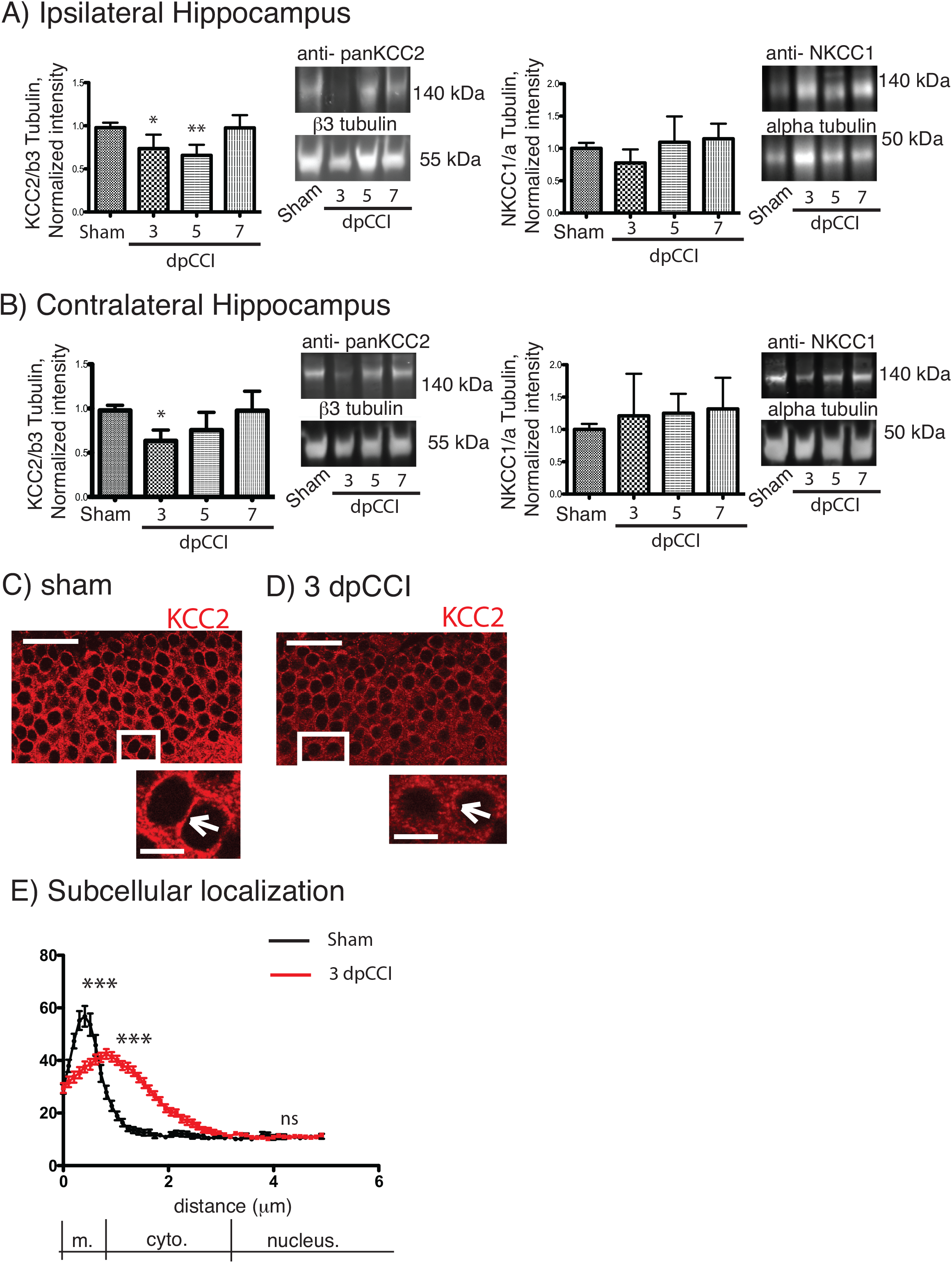
CCI induced changes in chloride co-transporters expression. A) The left panel represents the KCC2 protein expression normalized to the neuronal marker β3-tubulin on the ipsilateral hippocampus. Protein expression over the time is expressed in comparison to the sham conditions. On the right panel, NKCC1 protein expression is shown normalized to the ubiquitous marker α-tubulin. Protein expression over the time is expressed in comparison to sham conditions, n=8 per condition. One-way Anova test is performed and expressed as following *p<0,05; **p<0,01; ***p<0,001. B) same as A) but in the contralateral hippocampus. C-E KCC2 staining in granule cells. C) Sham at 3 dpCCI. The labelling is at the cellular membrane (arrowhead) and the cytoplasm is almost devoid of KCC2 labelling. D) 3 dpCCI. KCC2 is found in the cytoplasmic cell compartments (arrowheads). E) Histograms representing the distribution and quantification of the intensity of fluorescence in 90 3dpCCI cells (red curve) and 60 dpCCI cells in in sham (black curve). Statistical analysis represents the difference in each sub region of the cell, namely membrane, cytoplasm and nuclear staining. Scale bars: 50μm and 10μm.

### CCI induced internalization of KCC2 plasma membrane

We then investigate if residual KCC2 expression and more specifically its subcellular distribution was affected. We used a specific KCC2 antibody to examine the cellular distribution of KCC2. We decided to focus on DG which is an hippocampal region where changes in network activity is already reported after TBI (Bonislawski et al., 2007) and which is implicated in depression. In sham granular cells, KCC2 is mainly located near the membrane of cell bodies (Fig. 3D). In contrast, the labeling of KCC2 in granular cells is largely cytoplasmic 3dpCCI (Fig. 3E). The cellular distribution of KCC2 in sham and 3dpCCI is significantly different with a peak around the membrane for sham granular cells (sham 64,09 ± 4,550 N=2 n=60 vs 3dpCCI 44,34 ± 1,827 N=3 n=88) together with staining dispersion over the cytoplasmic compartment in granular cells at 3dpCCI (sham 13,56 ± 1,011 N=2 n=60 vs 3dpCCI 31,92 ± 1,538 N=3 n=88) (Fig. 3F). This suggests an internalization of KCC2 after TBI and is in agreement with robust changes in chloride homeostasis and GABAergic transmission in the DG.

### Bumetanide rescues post-traumatic impairment in secondary neurogenesis

Previous results have suggested that the effect of antidepressants on proliferation of adult born neurons of the DG may be involve in the mechanism of action of these compounds. Considering the prophylactic anti-depressant effect of bumetanide found in this study it is plausible that part of the antidepressive effect of bumetanide could be mediated by changes in proliferation. Thus, we monitored both the proliferative cells and newly born neurons at the end during the first post-traumatic week. Neurons were labelled with double-cortin (DCX), a marker of immature neurons (Ren et al., 2015) and proliferative cells were stained with bromo-deoxy-Uridin (BrdU) to assess the relative number of dividing cells within the granular layer of the DG (Samuels et al., 2015). In both cases the number of positive cells is calculated on a defined volume and expressed in raw data. We observed a significant CCI-induced reduction of the number of the DCX positive neurons within the DG both in the ipsi- and contralateral hippocampi at 7 dpCCI (sham 1 +/- 0.3 vs CCI ipsi 0.28 +/- 0.1*** and CCI contra 0.56 +/- 0.2**; respectively 6 and 4 animals, 3 to 4 slices per animal) (Fig. 4A and C) together with an increase in the number of BrdU positive cells within the contralateral DG (sham 1 +/- 0.3 vs CCI ipsi 1.18 +/- 0.1 and CCI contra 1.87 +/- 0.5***, n=4 and 6 animals 2 to 6 slices per animal) (Fig. 4A and C). Bumetanide treatment reduces first the number of BrdU positive cells (sham 1+/- 0.3 vs bum-CCI ipsi 0.6 +/- 0.2 and bum-CCI contra 1.2 +/- 0.3**) (Fig. 4A and C) and triggers an increase in the number of generated neurons (sham 1 +/- 0.2 bum-CCI ipsi 0.4 +/- 0.2 and bum-CCI contra 0.87 +/-0.2*) (Fig. 4A and C). Interestingly, one month after trauma, it was not possible to find any differences either in the number of BrdU+ or in the number of DCX+ neurons within the contralateral hippocampus of CCI animals (Fig. 4B and D). On the ipsilateral side at 1 month remains a significant and persistent loss of newly generated neurons with no significant change in BrdU positive cells (Fig. 4B and D) indicating a permanent change compared to the transient one observed in the contralateral side. Notably, bumetanide had no action on the ipsilateral hippocampus. This suggests a contribution of the contralateral hippocampus in the early settling up of post-traumatic depression.

**Figure 4:**
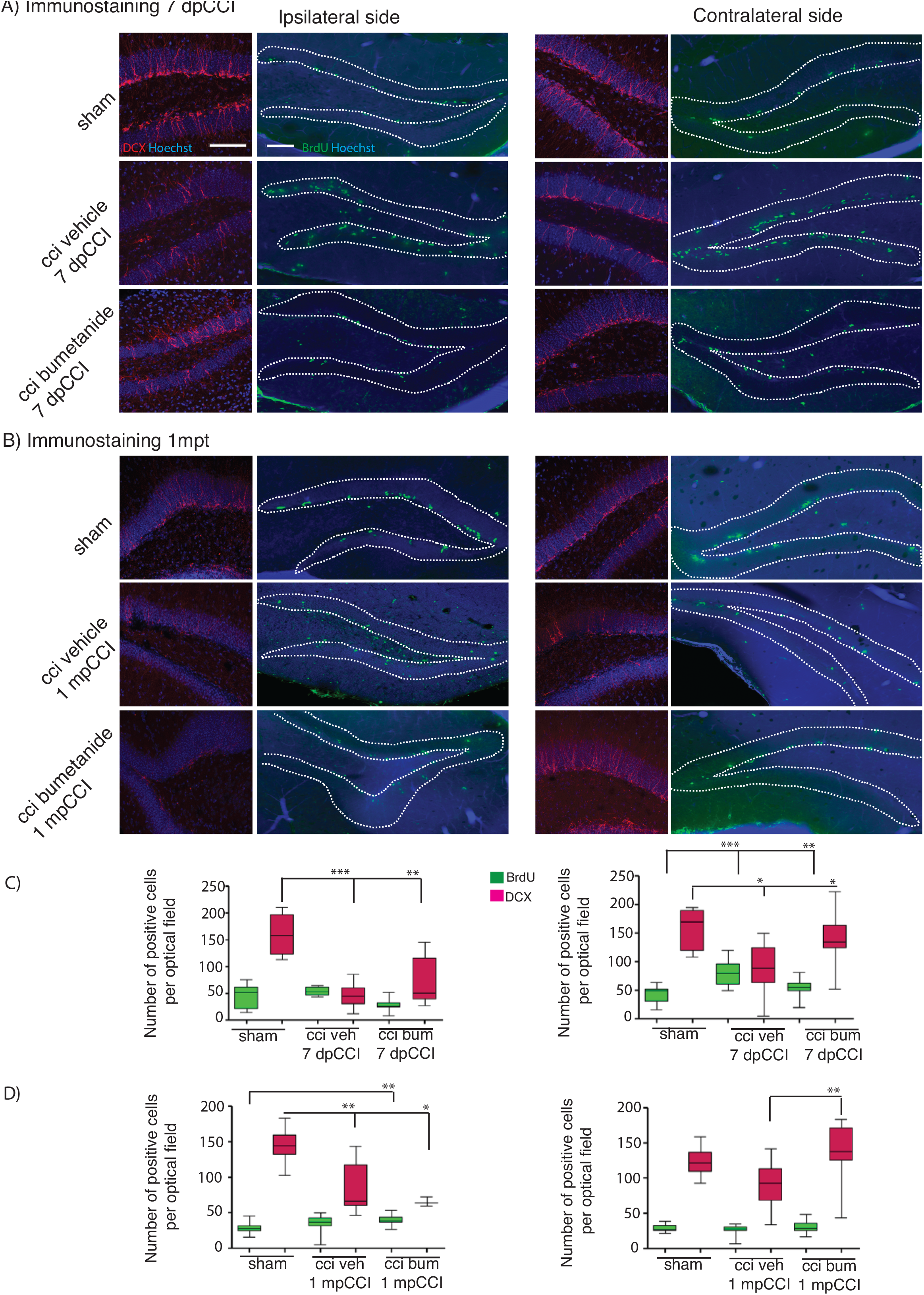
Effect of bumetanide on CCI induced changes in secondary neurogenesis. Secondary neurogenesis in dentate gyrus. A) Double-cortin (DCX) and BrdU labelling at 7 days post CCI in the ipsilateral (left) and contralateral (right) dentate gyrus of sham, CCI vehicle and bumetanide-treated animals. Dotted lines delimit granular layer of dentate gyrus (scale bar=100μm) B) Same as in (A) at 1 month post CCI. C) Quantification of BrdU and DCX positive cells 7 dpCCI in the ipsilateral (left) and contralateral (right) dentate gyrus of sham, CCI vehicle and bumetanide-treated animals. D) Same as in (C) at 1 month post CCI. DCX 7 days post CCI: n=6 animals per condition, 3 slices per animal; 1 month post CCI; n=4 animals per condition, 2-4 slices per animal. BrdU 7 days post CCI: n=5 sham n=6 CCI vehicle and 6 CCI bumetanide, 2-6 slices per animal, 1 month post CCI: n=3 sham n=4 CCI vehicle and 4 CCI bumetanide, 3-4 slices per animal.

Ambient GABA that is provided by the activity DG interneurons may play a role in the proliferation and migration of granular cell progenitors (Duan et al., 2008). We then wonder if GABAergic signaling known as a neurogenesis modulator (Alvarez et al., 2016; Moss and Toni, 2013; Samuels et al., 2015) could contribute to the etiology of post-traumatic depression (Cryan et al., 2005). Therefore, it appeared interesting to quantify parvalbumin interneurons, which are known to play critical role in post-traumatic consequences (Drexel et al., 2014; Hsieh et al., 2014; Khodaie et al., 2014). We quantified the parvalbumin-containing interneurons in the granular layer of the DG, both from ipsi and contralateral hippocampi at 7dpCCI. Both side showed a significant reduction of the number of parvalbumin-positive interneurons (ipsi 67% +/-23, ***, n=5 animals 25 slices; contra 40% +/-30, ***, n=5 animals 26 slices, Fig 5A, B). This loss was significantly reduced by bumetanide application at the contralateral (100% +/-31, ***, n=5 animals 3 to 4 slices per animal, Fig 5D) and ipsilateral side (52% +/-22, **, n=5 animals 3 to 4 slices per animal, Fig 5C) compared to CCI condition. Thus, the effect of bumetanide on DG secondary neurogenesis could be partly caused by changes in ambient GABA that is provided by the activity of DG interneurons.

**Figure 5:**
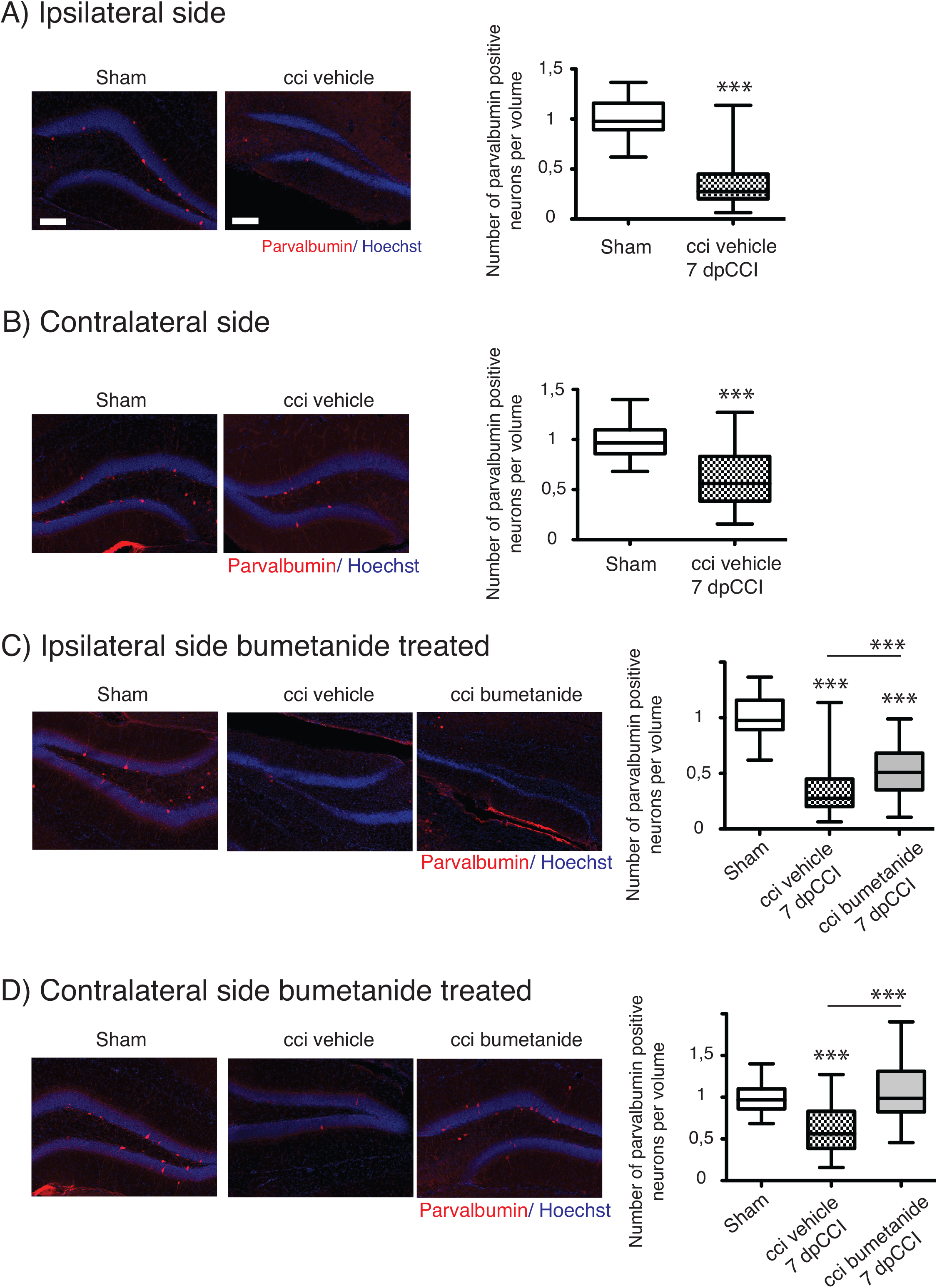
Effect of bumetanide on CCI induced parvalbumin positive interneuron death. A) Ipsilateral hippocampus: left panel example of parvalbumin and Hoechst immunostaining, from sham and CCI mice. On the right panel, quantification of parvalbumin positive interneurons in the dentate gyrus normalized to sham values. n = 5 animals per condition. B) Same as A) but on contralateral hippocampus, the histogram shows reduction in the number of parvalbumin-containing cells in the DG, n = 5 animals per condition. C) Effect of bumetanide in parvalbuming interneuron survival in the ipsilateral hippocampus. The histogram shows a significant reduction in the cell loss in the presence of bumatanide but this is though significantly less as compared to compared to sham, n = 5 animals per condition. D) Contralateral hippocampus: bumetanide injection reduces interneurons loss, n = 5 animals per condition. One-way Anova test is expressed as following *p<0,05; **p<0,01; ***p<0,001.

## Discussion

Here we found that CCI-induced DLB is strongly sensitive to trauma-induced changes in GABA(A) mediated responses. The block of depolarizing GABA(A) responses by bumetanide, at very early stages after CCI, significantly leads to amelioration DLB at chronic stages. The question raised by these results is whether and how changes in GABA(A) transmission are involved in the CCI-induced rearrangements in hippocampus leading to abnormal behavior.

Our behavioral analyses after CCI highlight results on long-term changes indicating impairment of mood associated behavior. CCI mice exhibited a phenotype that mimick a decreased defensive behavior in an unfamiliar environment, as it has been descrived in other anxiety-like behavior tests (Pandey et al., 2009; Stemper et al., 2015). Surprisingly, blocking GABA(A) mediated depolarization with the specific inhibitor bumetanide at early stages after CCI resulted in a long-term significant reduction in DLB, long after the end of the treatment with bumetanide. These results pinpoint an important role of the qualitative changes in GABA(A) responses and suggests that bumetanide itself could act in a prophylactic manner as anti-depressant compound, this was proven to be independent on its effect on lesion volume in a model of cerebral ischemia (Xu et al., 2016).

Consistent changes in KCC2 and NKCC1 expression have been found in a number of trauma models as well as in resections form temporal lobe epilepsy (Pallud et al., 2014). Qualitative changes in GABA(A) transmission and levels of chloride regulatory proteins have however not been well characterized in CCI models (Hui et al., 2016; Robel et al., 2015). In the present study we showed that KCC2 but not NKCC1 expression is significantly changed in the hippocampus during the first week following trauma. Interestingly, although these changes occur shortly after CCI in both the ipsi and contralateral hippocampus they are transient as KCC2 expression levels return to normal values in both hippocampi. Changes in chloride extrusion efficacy are consistent with this biochemical conclusion. Thus resulting in a switch in GABA(A) mediated network excitability as opposed to control condition. As expected the effects observed in the contralateral hippocampus are milder but present.

Previous results have shown that almost all antidepressants have significant effects on proliferation in the DG and production of newborn neurons. This has led to the hypothesis that proliferation in the DG is associated to DLB. While CCI induced significant increase proliferation only in the contralateral DG the number of DCX positive immature neurons were significantly diminished on both sides. The acute effect of bumetanide was to increase neurons production and to reduce the proliferation. Although on the ipsilateral side these effect were not longer present after one month, the contralateral side showed significant increase number of DCX positive cell in CCI mice compared to bumetanide-treated. These results provide evidences that transition from BrdU positive cells to double-cortin positive immature neurons after CCI is significantly affected by blocking GABA(A) mediated responses at early stages and that this effect outlast significantly after the end of bumetanide application affecting both the functional changes in chloride transporters and the qualitative GABA(A) responses.

Although the CCI-induced changes in chloride regulatory proteins and GABA(A) transmission are consistent with short-term effects on excitability. It is less obvious how this is involved in long-term effect after trauma. A possible contributing explanation could be related to cell death of interneurons. We have previously shown that the qualitative changes in GABA(A) responses is tightly linked with the mechanism for survival of injured neurons (Pellegrino et al., 2011; Shulga et al., 2012). Changes in interneurons population leading to changes in GABA release could significantly change the excitability of the network (Hsieh et al., 2014; Shiri et al., 2014). In some pathological contexts such as temporal lobe epilepsy and TBI as well as in other different model of acquired epilepsy, parvalbumin interneurons are known to be very sensitive to death (Drexel et al., 2011; Hsieh et al., 2014). We and others have proposed that bumetanide could prevent trauma induced cell death (Hui et al., 2016; Shulga and Rivera, 2013). This mechanism involved the block of post traumatic depolarizing effect of GABA(A) receptor that is produced by KCC2 functional down regulation (Hui et al., 2016; Lee et al., 2011; Pellegrino et al., 2011; Winkelmann et al., 2015). The results presented here clearly show that bumetanide prevents CCI-induced interneuron death at least in the DG. Thus, the previously shown mechanism for trauma-triggered apoptosis of principal cells could be also relevant for interneurons. The activity of parvalbumin interneurons has been linked to changes in secondary neurogenesis in the DG. Released ambient GABA form hilar interneurons is of primary importance for regulating the proliferation state of cells within the DG (Alvarez et al., 2016; Boldrini et al., 2013; Moss and Toni, 2013; Samuels et al., 2015). Thus, the effect of bumetanide on interneuron survival could contribute to the short and long-term effects on DG proliferation (Sun et al., 2012).

In the present study we have manly focused on DLB but bumetanide amelioration of other pathological behavior may be also present. It will be highly interesting to supplement these studies with others cognitive tests for e.g. learning and memory together with social interaction paradigms. We showed that each hemisphere react to the brain trauma in different extend and with different kinetic. This highlight that the consequences in the contralateral hemisphere are as important as the ipsilateral side (Khalilov et al., 2003) and thus cannot be considered as an independent structure in the etiology of DLB. This results also prompts testing specific compound to block NKCC1 that are able to penetrate the blood brain barrier or promote KCC2 function after CCI to restore GABAergic activity (Gagnon et al., 2013; Medina et al., 2014; Puskarjov et al., 2014).

In conclusion the results presented here present a novel mechanism for involved in the development of TBI induced DLB and may open new interesting perspectives to treat TBI associated psychiatric conditions and suggests the used of bumetanide as an interesting prophylactic agent.

## Acknowledgements

We are grateful to Pr.Y. Ben-Ari and Drs. G. Chazal & F. Molinari for critical reading of the manuscript. Some data in this article are part of the doctoral thesis of M.A. presented to the Medical Faculty of the Johannes Gutenberg-University Mainz, Germany. This work is supported by the French national agency for research, ANR through CR and EG ANR-Traumep-13-BSV4-0012-01, the Eranet Neuron III program through Acrobat grant to CR, the Akademy of Finland (AK1308265) to CR and by the “fondation des gueules cassées” through EG, part of the behavioral unit has been funded by an FRC grant “espoir en tête 2015” to Y. Ben-Ari.

---

**Supplemental 1.**
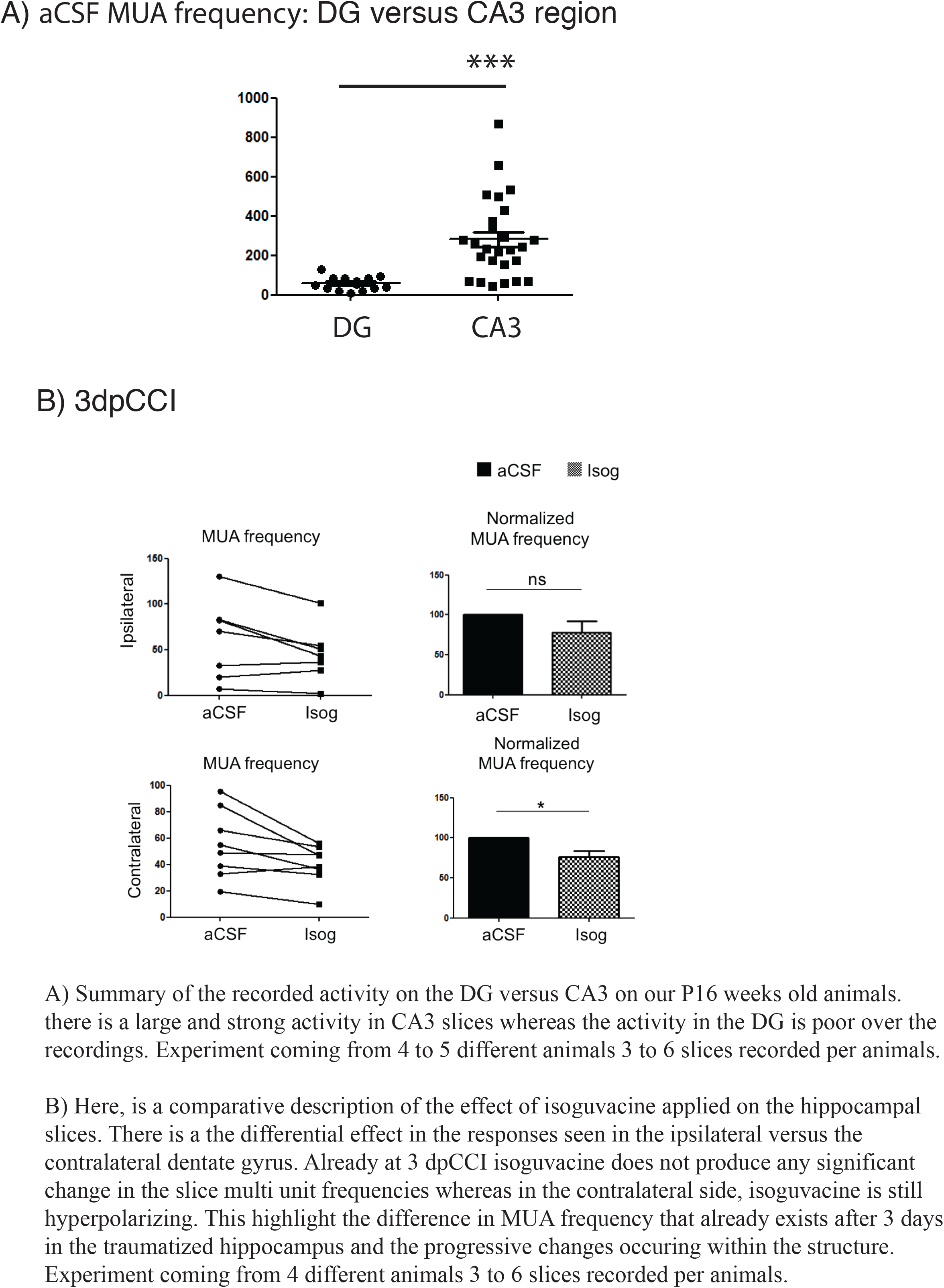
A) Summary of the recorded activity on the DG versus CA3 on our P16 weeks old animals. there is a large and strong activity in CA3 slices whereas the activity in the DG is poor over the recordings. Experiment coming from 4 to 5 different animals 3 to 6 slices recorded per animals. B) Here, is a comparative description of the effect of isoguvacine applied on the hippocampal slices. There is a the differential effect in the responses seen in the ipsilateral versus the contralateral dentate gyrus. Already at 3 dpCCI isoguvacine does not produce any significant change in the slice multi unit frequencies whereas in the contralateral side, isoguvacine is still hyperpolarizing. This highlight the difference in MUA frequency that already exists after 3 days in the traumatized hippocampus and the progressive changes occuring within the structure. Experiment coming from 4 different animals 3 to 6 slices recorded per animals.

**Supplemental figure 2.**
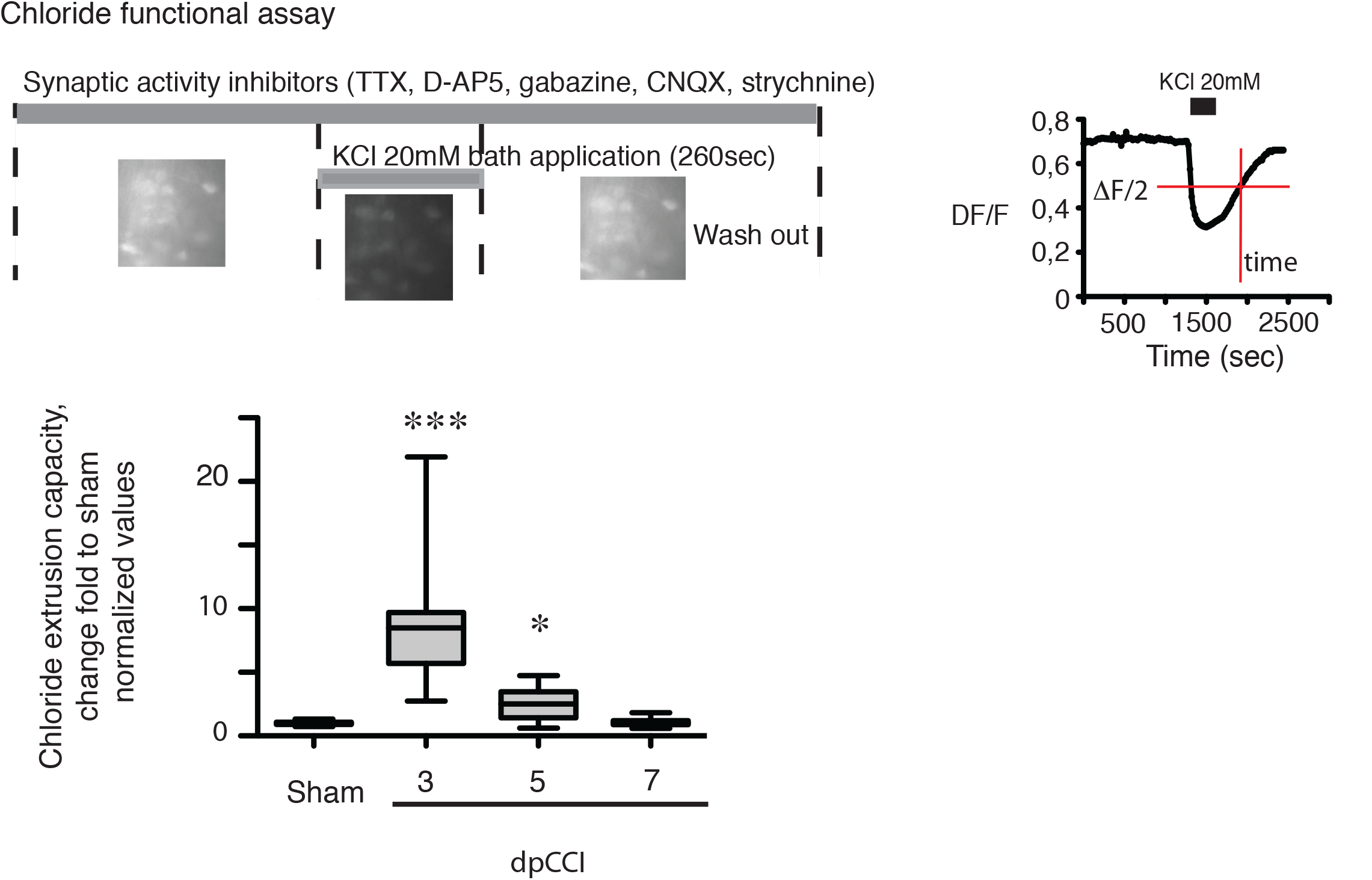
Experimental design of the chloride recording: Bath application of 25mM potassium chloride solution for 260sec in presence of all synaptic blockers. Both the amplitude of the response triggered by the bath application and the recovery of the signal is quantified. Results are represented as chloride extrusion capacity by reporting the fluorescence recovery, meaning the extrusion capacity depending on the amplitude of the signal, to avoid any bias due to changes in loading amplitude.

**Supplemental 3.**
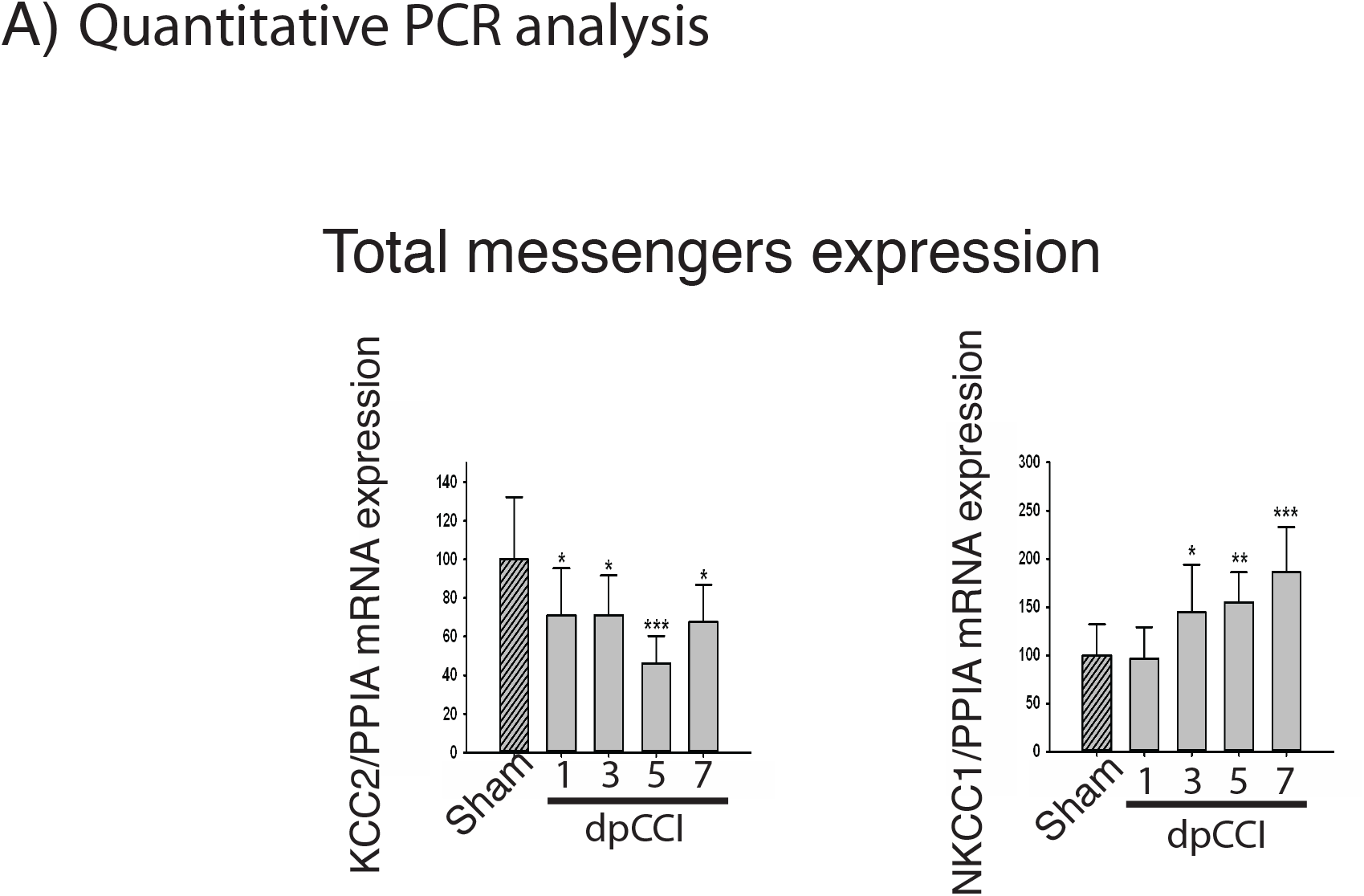
A) Relative KCC2 and NKCC1 mRNA expression is normalized to cyclophilin A gene (PPIA) at different time after trauma during the first post-traumatic week (n=10 per condition). One-way Anova test is performed and expressed as following *p<0,05; **p<0,01; ***p<0,001.

